# PCP4 immunoreactivity suggests the presence of hippocampal region CA2 in solitary, social and eusocial mole-rat species

**DOI:** 10.1101/2023.02.02.526898

**Authors:** Tristan M. Stöber, Maria K. Oosthuizen

## Abstract

Because African mole-rats express diverse social behaviors, they are prime candidates to study the effect of sociability on the evolution of brain circuits. This study compared the expression of Purkinje Cell Protein 4 (PCP4) in hippocampal slices of solitary Cape mole-rats, social Highveld mole-rats, and eusocial Damaraland and naked mole-rats. PCP4 is an established marker for pyramidal cells of hippocampal region CA2, a key structure for social memory and aggression. We observed prominent PCP4 immunoreactivity in the central part of the *cornu Ammonis* for all mole-rat species. While further verification is required, these findings suggest the extend of CA2 does not drastically differ despite varying social behaviors. Future studies may use this work as a starting point to explore the relationship between environmental requirements and the functional role of CA2.

## 1 Introduction

African mole-rats (family Bathyergidae) are subterranean rodents that are endemic to sub-Saharan Africa. This family is remarkable in several ways. Spending their lives almost entirely in extensive underground tunnel systems, they have attracted attention due to their long life expectancy (Edrey et al., 2012, 2013; Dammann et al., 2011), resistance to tumors (Liang et al., 2010), adaptations to low oxygen concentrations (Peterson et al., 2012; Park et al., 2017), and diverse social behaviors, ranging from strictly solitary to social and eusocial species.

This work considers four mole-rat species covering the full spectrum of sociality: The strictly solitary Cape mole-rat (*Georychus capensis*), the social Highveld mole-rat (*Cryptomys hottentotus*), as well as the eusocial Damaraland mole-rat (*Fukomys damarensis*) and naked mole-rat (*Heterocephalus glaber*). While Cape mole-rats meet each other with deadly aggression outside brief periods for mating and maternal care (Bennett and Jarvis, 1988), Highveld mole-rats form colonies with one breeding pair and up to 12 subordinates (Bennett and Faulkes, 2000). Damaraland mole-rat and naked mole-rats live in larger and more complex colonies with social hierarchies, division of labor, and cooperative breeding (Jarvis, 1981; Jarvis and Bennett, 1993). Thus, due to their evolutionary proximity, similar ecological niche, and life history, these species are prime candidates for comparative studies to understand the effect of sociality on brain circuit evolution (Kalamatianos et al., 2010; Amrein et al., 2014; Coen et al., 2015; Kverková et al., 2018).

Recent experimental findings established the CA2 region of the hippocampus as a key structure for social recognition memory and aggression (Hitti and Siegelbaum, 2014; Stevenson and Caldwell, 2014; Smith et al., 2016; Piskorowski et al., 2016; Leroy et al., 2017; Meira et al., 2018; Okuyama et al., 2016; Dominguez et al., 2019; Oliva et al., 2020; Donegan et al., 2020; Leroy et al., 2021; Boyle et al., 2022; Loisy et al., 2022, but see also Lehr et al., 2021 for its broader role in hippocampal information processing). Without dorsal CA2, mice appear not to remember that they have already interacted with a conspecific (Hitti and Siegelbaum, 2014; Stevenson and Caldwell, 2014) and are less likely to aggressively attack an intruder mouse (Leroy et al., 2018). Both recognition and aggression depend on the vasopressin receptor 1b (Wersinger et al., 2002; Leroy et al., 2018), which is expressed almost exclusively in the dorsal CA2 (Young et al., 2006).

The importance of CA2 for social behavior motivated us to investigate the prominence of the hippocampal region CA2 in solitary, social and eusocial mole-rat species. As a first step, we characterized the CA2 region by staining for Purkinje Cell Protein 4 (PCP4). PCP4 is one of several established molecular markers for CA2 pyramidal cells in mice (San Antonio et al., 2014; Kohara et al., 2014). We observed prominent PCP4 immunoreactivity in the central part of the *cornu Ammonis* for all mole-rat species. These data suggest that the extend of CA2 does not drastically deviate despite extreme differences in social behavior. However, given the superficial nature of this analysis, further verification is warranted.

## 2 Methods

### 2.1 Animals

We used remnant mole-rat slices from two previous studies (Amrein et al., 2014, Highveld, Cape and naked mole-rats) and (Oosthuizen and Amrein, 2016, Damaraland mole-rats). Each mole-rat species had a different origin: Cape mole-rats were trapped near Darling, Western Cape, South Africa, and Highveld mole rats in Centurion, Gauteng, South Africa. Damaraland mole-rats originated from Blackrock, Northern Cape, South Africa. Naked mole-rats were derived from a colony at the University of Nairobi, Kenya. Trapping and treatment were in accordance with the ethical guidelines of the respective universities. Neither queens nor pregnant or lactating animals were included in the study. After weighing and determining sex, the animals were anesthetized with pentobarbital, 50 mg / kg body weight, and perfused transcardiacally with phosphate buffered saline (PBS), followed by sodium sulfide solution and fixed with 4% paraformaldehyde (PFA). The brains were extracted, post-fixed overnight and divided into the right and left hemispheres, which were cryoprotected in 30% sucrose and processed for immuno-histochemistry, or stored in PFA until methacrylate embedding, respectively. Teeth and femurs were extracted for subsequent age determination.

### 2.2 Histology

To stain for PCP4 expression, we used Rabbit anti-PCP4, HPA005792, BIOZOL Diagnostic, lot number 15962 and followed this protocol, with free floating sections and constant agitation: a) Washed 4×10 min in PBS-T (PBS + 0.5% Triton X-100), b) incubated in antibody diluent (PBS-T + 10% normal goat serum NGS) for 1 hour, c) incubated with primary antibody anti-PCP4 1:200 in PBS-T + 5% NGS over night, d) rinsed 3×5 min in PBS-T, e) incubated with secondary antibody (goat anti-rabbit Alexa 488) 1:200 in PBS for 2 hours, f) 1:15000 DAPI in PBS 1X5min (also for positive and negative controls), g) rinsed 5×5 minutes in PBS, h) mounted, rinsed in ddH2O, and coverslipped with antifade Fluorogold. Steps e) to h) were performed in darkness.

## 3 Results

To characterize CA2 in mole-rat species throughout the full spectrum of sociality, we investigated PCP4 expression in hippocampal slices of strictly solitary Cape mole-rat, the social Highveld mole-rat, as well as the eusocial Damaraland mole-rat and naked mole-rat. We detected spatially restricted PCP4 expression in a central part of the *cornu Ammonis* pyramidal cell layer in all four mole-rat species, similar to the common laboratory mouse strain *C57BL/6* (see Fig. 1). These findings suggest the presence of CA2 in all investigated mole-rat species.

**Figure 1:**
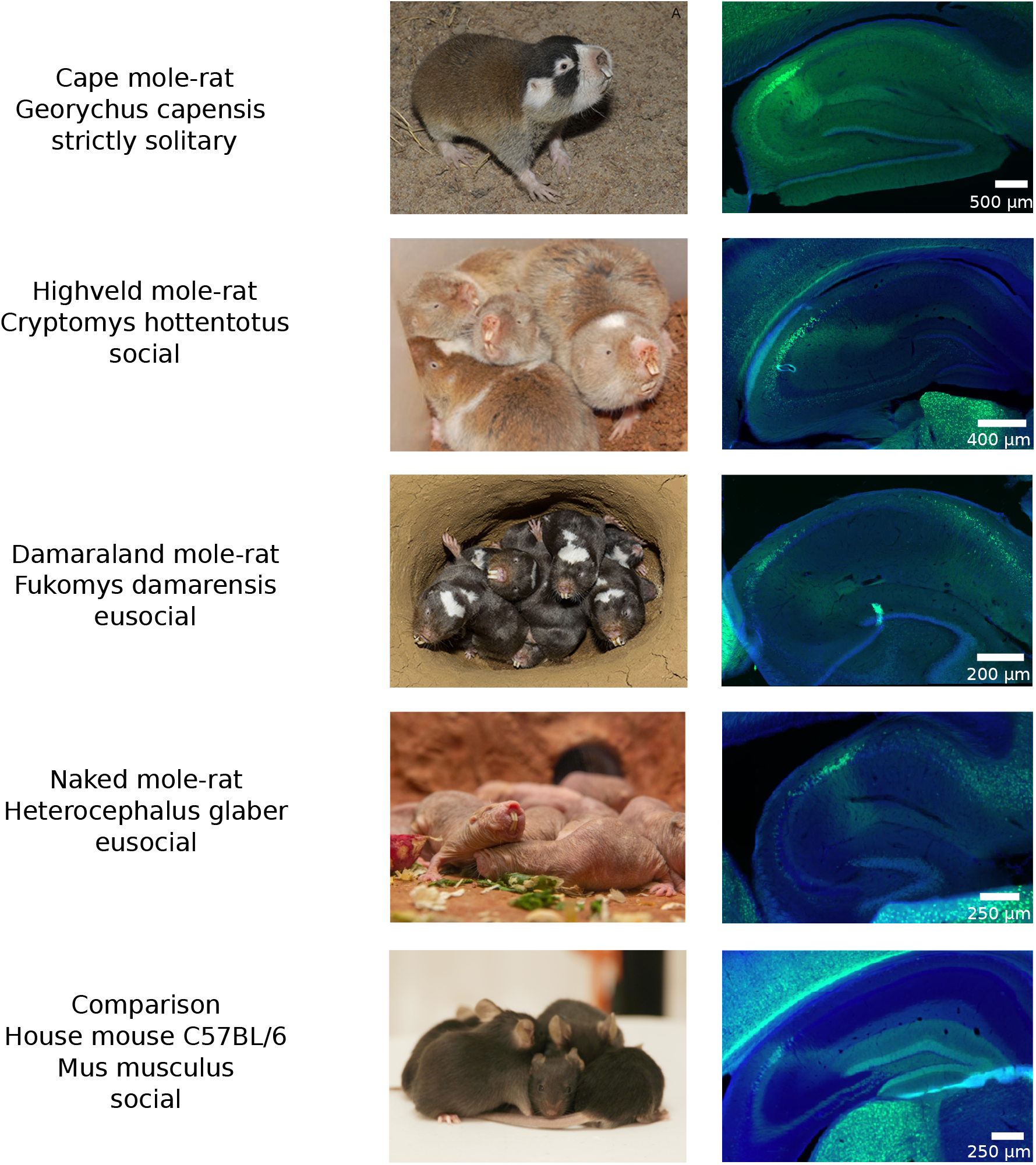
PCP4 immunoreactivity in a central region of the *cornu Ammonis* suggests the presence of hippocampal region CA2 in solitary, social and eusocial mole-rats species. Note, the immunostaining in the Highveld mole-rat has been performed in a different subspecies (*Cryptomys hottentotus pretoriae*) compared to the picture depicting a group of animals (*Cryptomys hottentotus hottentotus*). Pictures of living animals are adapted from the following sourses: Cape and Damaraland mole-rat Oosthuizen (2017), Highveld mole-rat Elizabeth Archer (personal communication), naked mole-rat Smithsonian’s National Zoological Park shared under CC0, house mouse by Boris Heifets shared under CC BY 4.0.

## 4 Discussion

In this study, we used PCP4 immunostaining to label hippocampal region CA2 in hippocampal slices of four different mole-rat species with diverse levels of sociability and compared them to the common house mouse. In all species, we found prominent staining in a central part of the *cornu Ammonis* region - which indicates the presence of CA2. However, to corroborate these findings future work should combine staining with multiple CA2 specific markers, such as RGS14, STEP, and *α* -actinin 2 (Lee et al., 2010; Kohara et al., 2014; Chevaleyre and Siegelbaum, 2010), and classical anatomical markers, such as the presence of pyramidal cells with mossy fiber input but without complex spines (Kohara et al., 2014).

Given the broad range of sociality in mole-rats, differences in CA2 would not come as a surprise. Previous investigations found that neurogenesis is generally low in mole-rats, but the amount of neurogenesis in the dentate gyrus differs between social and solitary species. Proportions of young neurons are lower in solitary Cape mole-rats, compared to social Highveld and naked mole-rats (Amrein et al., 2014). Furthermore, in both eusocial naked and Damaraland mole-rats an inverse relationship between social status and neurogenesis has been observed, with queens showing the least and workers the most neurogenesis (Peragine et al., 2014; Oosthuizen and Amrein, 2016). It seems plausible that the observed differences in neurogenesis reflect distinct cognitive requirements, both in habitat and social interactions (Oosthuizen, 2017), as has been observed in rats (Kozorovitskiy and Gould, 2004) and mice (Monteiro et al., 2014). Along similar lines, it can be argued that requirements for social recognition memory likely differ in solitary and social mole-rat species and, thus, should properties of CA2.

A thorough anatomical description of CA2 in mole-rats would open new research avenues. An area of interest may be the potential relationship between species-specific aggression levels and differences in CA2. Mice exhibit social aggression mediated by the activity of pyramidal cells in dorsal CA2 (Wersinger et al., 2002; Leroy et al., 2018). As mice transition from exploratory behavior to aggression, the firing rates of these pyramidal cells gradually increase, activating the lateral septum and subsequently disinhibiting the ventromedial hypothalamus, which is known to initiate an attack (Leroy et al., 2018). The function of this circuit depends on the CA2-specific vasopressin receptor 1b. Infusing an antagonist of the vasopressin receptor 1b has been shown to prevent aggression in mice (Leroy et al., 2018). Thus, it can be hypothesized that in Cape mole-rats the suppression of aggression during mating and maternal care may be regulated by vasopressin receptors in CA2. Using a similar antagonist infusion as in mice, this hypothesis can be directly tested.

## 5 Acknowledgements

We thank 1) Jovana Maliković and Irmgard Amrein, as well as the Center of Microscopy and Image Analysis, University of Zurich, Switzerland, for the PCP4 immunostaining, 2) Nilesh B. Patel from the University of Nairobi, Kenya, for providing brain tissue of naked mole-rats, 3) Elizabeth Archer for providing the picture of a group of Highveld mole-rats, 4) Boris Heifets for providing a picture of a group of laboratory mice, 5) Ane Charlotte Christensen for providing the PCP4 staining protocol.

## References

Amrein, I., Becker, A. S., Engler, S., Huang, S.-h., Müller, J., Slomianka, L., and Oosthuizen, M. K. (2014). Adult neurogenesis and its anatomical context in the hippocampus of three mole-rat species. Frontiers in Neuroanatomy, 8:39.

Bennett, N. and Jarvis, J. (1988). The reproductive biology of the cape mole-rat, georychus capensis (rodentia, bathyergidae). Journal of Zoology, 214(1):95–106.

Bennett, N. C. and Faulkes, C. G. (2000). African mole-rats: Ecology and Eusociality. Cambridge University Press.

Boyle, L., Posani, L., Irfan, S., Siegelbaum, S. A., and Fusi, S. (2022). The geometry of hippocampal CA2 representations enables abstract coding of social familiarity and identity. bioRxiv.

Chevaleyre, V. and Siegelbaum, S. A. (2010). Strong CA2 pyramidal neuron synapses define a powerful disynaptic cortico-hippocampal loop. Neuron, 66(4):560–572.

Coen, C. W., Kalamatianos, T., Oosthuizen, M. K., Poorun, R., Faulkes, C. G., and Bennett, N. C. (2015). Sociality and the telencephalic distribution of corticotrophin-releasing factor, urocortin 3, and binding sites for crf type 1 and type 2 receptors: A comparative study of eusocial naked mole-rats and solitary cape mole-rats. Journal of Comparative Neurology, 523(16):2344–2371.

Dammann, P., Šumbera, R., Maßmann, C., Scherag, A., and Burda, H. (2011). Extended longevity of reproductives appears to be common in fukomys mole-rats (rodentia, bathyergidae). PLoS One, 6(4):e18757.

Dominguez, S., Rey, C. C., Therreau, L., Fanton, A., Massotte, D., Verret, L., Piskorowski, R. A., and Chevaleyre, V. (2019). Maturation of PNN and ErbB4 signaling in area CA2 during adolescence underlies the emergence of PV-interneuron plasticity and social memory. Cell Reports, 29(5):1099–1112.

Donegan, M. L., Stefanini, F., Meira, T., Gordon, J. A., Fusi, S., and Siegelbaum, S. A. (2020). Coding of social novelty in the hippocampal CA2 region and its disruption and rescue in a 22q11.2 microdeletion mouse model. Nature Neuroscience, 23(11):1365–1375.

Edrey, Y. H., Casper, D., Huchon, D., Mele, J., Gelfond, J. A., Kristan, D. M., Nevo, E., and Buffenstein, R. (2012). Sustained high levels of neuregulin-1 in the longest-lived rodents; a key determinant of rodent longevity. Aging cell, 11(2):213–222.

Edrey, Y. H., Medina, D. X., Gaczynska, M., Osmulski, P. A., Oddo, S., Caccamo, A., and Buffenstein, R. (2013). Amyloid beta and the longest-lived rodent: the naked mole-rat as a model for natural protection from alzheimer’s disease. Neurobiology of aging, 34(10):2352–2360.

Hitti, F. L. and Siegelbaum, S. A. (2014). The hippocampal CA2 region is essential for social memory. Nature, 508(7494):88–92.

Jarvis, J. (1981). Eusociality in a mammal: cooperative breeding in naked mole-rat colonies. Science, 212(4494):571–573.

Jarvis, J. and Bennett, N. (1993). Eusociality has evolved independently in two genera of bathyergid mole-rats—but occurs in no other subterranean mammal. Behavioral Ecology and Sociobiology, 33(4):253–260.

Kalamatianos, T., Faulkes, C. G., Oosthuizen, M. K., Poorun, R., Bennett, N. C., and Coen, C. W. (2010). Telencephalic binding sites for oxytocin and social organization: A comparative study of eusocial naked mole-rats and solitary cape mole-rats. Journal of Comparative Neurology, 518(10):1792–1813.

Kohara, K., Pignatelli, M., Rivest, A. J., Jung, H.-Y., Kitamura, T., Suh, J., Frank, D., Kajikawa, K., Mise, N., Obata, Y., et al. (2014). Cell type-specific genetic and optogenetic tools reveal hippocampal CA2 circuits. Nature Neuroscience, 17(2):269–279.

Kozorovitskiy, Y. and Gould, E. (2004). Dominance hierarchy influences adult neurogenesis in the dentate gyrus. Journal of Neuroscience, 24(30):6755–6759.

Kverková, K., Bělíková, T., Olkowicz, S., Pavelková, Z., O’Riain, M. J., Šumbera, R., Burda, H., Bennett, N. C., and Němec, P. (2018). Sociality does not drive the evolution of large brains in eusocial african mole-rats. Scientific reports, 8(1):1–14.

Lee, S. E., Simons, S. B., Heldt, S. A., Zhao, M., Schroeder, J. P., Vellano, C. P., Cowan, D. P., Ramineni, S., Yates, C. K., Feng, Y., et al. (2010). RGS14 is a natural suppressor of both synaptic plasticity in CA2 neurons and hippocampal-based learning and memory. Proceedings of the National Academy of Sciences, 107(39):16994–16998.

Lehr, A. B., Kumar, A., Tetzlaff, C., Hafting, T., Fyhn, M., and Stöber, T. M. (2021). CA2 beyond social memory: Evidence for a fundamental role in hippocampal information processing. Neuroscience & Biobehavioral Reviews, 126:398–412.

Leroy, F., Brann, D. H., Meira, T., and Siegelbaum, S. A. (2017). Input-timing-dependent plasticity in the hippocampal CA2 region and its potential role in social memory. Neuron, 95(5):1089–1102.

Leroy, F., de Solis, C. A., Boyle, L. M., Bock, T., Lofaro, O. M., Buss, E. W., Asok, A., Kandel, E. R., and Siegelbaum, S. A. (2021). Enkephalin release from vip interneurons in the hippocampal ca2/3a region mediates heterosynaptic plasticity and social memory. Molecular Psychiatry, pages 1–22.

Leroy, F., Park, J., Asok, A., Brann, D. H., Meira, T., Boyle, L. M., Buss, E. W., Kandel, E. R., and Siegelbaum, S. A. (2018). A circuit from hippocampal ca2 to lateral septum disinhibits social aggression. Nature, 564(7735):213–218.

Liang, S., Mele, J., Wu, Y., Buffenstein, R., and Hornsby, P. J. (2010). Resistance to experimental tumorigenesis in cells of a long-lived mammal, the naked mole-rat (heterocephalus glaber). Aging cell, 9(4):626–635.

Loisy, M., Bouisset, G., Lopez, S., Muller, M., Spitsyn, A., Duval, J., Piskorowski, R. A., Verret, L., and Cheva-leyre, V. (2022). Sequential inhibitory plasticities in hippocampal area ca2 and social memory formation. Neuron.

Meira, T., Leroy, F., Buss, E. W., Oliva, A., Park, J., and Siegelbaum, S. A. (2018). A hippocampal circuit linking dorsal CA2 to ventral CA1 critical for social memory dynamics. Nature Communications, 9(1):4163.

Monteiro, B. M., Moreira, F. A., Massensini, A. R., Moraes, M. F., and Pereira, G. S. (2014). Enriched environment increases neurogenesis and improves social memory persistence in socially isolated adult mice. Hippocampus, 24(2):239–248.

Okuyama, T., Kitamura, T., Roy, D. S., Itohara, S., and Tonegawa, S. (2016). Ventral CA1 neurons store social memory. Science, 353(6307):1536–1541.

Oliva, A., Fernández-Ruiz, A., Leroy, F., and Siegelbaum, S. A. (2020). Hippocampal CA2 sharp-wave ripples reactivate and promote social memory. Nature, 587(7833):264–269.

Oosthuizen, M. K. (2017). From mice to mole-rats: species-specific modulation of adult hippocampal neurogenesis. Frontiers in neuroscience, 11:602.

Oosthuizen, M. K. and Amrein, I. (2016). Trading new neurons for status: Adult hippocampal neurogenesis in eusocial damaraland mole-rats. Neuroscience, 324:227–237.

Park, T. J., Reznick, J., Peterson, B. L., Blass, G., Omerbašić, D., Bennett, N. C., Kuich, P. H. J., Zasada, C., Browe, B. M., Hamann, W., et al. (2017). Fructose-driven glycolysis supports anoxia resistance in the naked mole-rat. Science, 356(6335):307–311.

Peragine, D., Simpson, J., Mooney, S., Lovern, M., and Holmes, M. (2014). Social regulation of adult neurogenesis in a eusocial mammal. Neuroscience, 268:10–20.

Peterson, B. L., Larson, J., Buffenstein, R., Park, T. J., and Fall, C. P. (2012). Blunted neuronal calcium response to hypoxia in naked mole-rat hippocampus. PloS one, 7(2):e31568.

Piskorowski, R. A., Nasrallah, K., Diamantopoulou, A., Mukai, J., Hassan, S. I., Siegelbaum, S. A., Gogos, J. A., and Chevaleyre, V. (2016). Age-dependent specific changes in area CA2 of the hippocampus and social memory deficit in a mouse model of the 22q11.2 deletion syndrome. Neuron, 89(1):163–176.

San Antonio, A., Liban, K., Ikrar, T., Tsyganovskiy, E., and Xu, X. (2014). Distinct physiological and developmental properties of hippocampal ca2 subfield revealed by using anti-purkinje cell protein 4 (pcp4) immunostaining. Journal of Comparative Neurology, 522(6):1333–1354.

Smith, A., Avram, S. W., Cymerblit-Sabba, A., Song, J., and Young, W. (2016). Targeted activation of the hippocampal CA2 area strongly enhances social memory. Molecular Psychiatry, 21(8):1137–1144.

Stevenson, E. L. and Caldwell, H. K. (2014). Lesions to the CA2 region of the hippocampus impair social memory in mice. European Journal of Neuroscience, 40(9):3294–3301.

Wersinger, S., Ginns, E. I., O’carroll, A., Lolait, S., and Young, W. (2002). Vasopressin V1b receptor knockout reduces aggressive behavior in male mice. Molecular Psychiatry, 7(9):975–984.

Young, W. S., Li, J., Wersinger, S. R., and Palkovits, M. (2006). The vasopressin 1b receptor is prominent in the hippocampal area CA2 where it is unaffected by restraint stress or adrenalectomy. Neuroscience, 143(4):1031–1039.

